# Genetic draft and valley crossing

**DOI:** 10.1101/383737

**Authors:** Taylor Kessinger, Jeremy Van Cleve

## Abstract

Living systems are characterized by complex adaptations which require multiple coordinated mutations in order to function. Empirical studies of fitness landscapes that result from the many possible mutations in a gene region reveal many fitness peaks and valleys that connect them. Thus, it is possible that some complex adaptations have arisen by evolutionary paths whose intermediate states are neutral or even deleterious. When intermediates are deleterious, traversing such an evolutionary path is known as “crossing a fitness valley”. Previous efforts at studying this problem have rigorously characterized the rate at which such complex adaptations evolve in populations of roughly equally fit individuals. However, populations that are very large or have broad fitness distributions, such as many microbial populations, adapt quickly, which substantially alters the fate and dynamics of individual mutations due to the action of genetic draft. We investigate the rate at which complex adaptations evolve in these rapidly adapting populations in regions without recombination. We confirm that rapid adaptation overall increases the time required to cross a valley; however, rapid adaptation can make it easier for deeper valleys to be crossed relative to the time required for single beneficial mutations to sweep to fixation.

## Introduction

The simplest adaptive scenarios in evolution involve the arisal and fixation of successive beneficial mutations. This appears to be how Darwin thought even complex systems, such as the vertebrate eye, evolved (Darwin 1859), and this assumption underlies many models in evolutionary theory including those from adaptive dynamics (Geritz *et al*. 1998; Dercole and Rinaldi 2008) and theories of adaptation (Gillespie 1983; Orr 1998; Gillespie 1991). Evolution will certainly proceed in this fashion if there always exists a sequence of mutations from an initial genotype to the high fitness genotype in which each mutant genotype in the sequence has a higher fitness than the previous one. If the individual fitness of each genotype is visualized as a surface or landscape where the axes represent alternative alleles at each locus, then smooth landscapes with a single peak ensure that uphill paths exist no matter where a population starts. However, many empirical fitness landscapes are not completely smooth and have multiple peaks (reviewed in Szendro *et al*. 2013; de Visser and Krug 2014; Obolski *et al*. 2018). In such landscapes, wild type individuals may have to traverse a mutational valley–a region of lower fitness–in order to reach a higher fitness peak. This is a special case of “sign epistasis”. We seek to characterize this “valley crossing” process in order to better understand the routes adaptation is likely to take on these rugged empirical landscapes (e.g., Aguilar-Rodríguez *et al*. 2017) and to assess the likelihood of valley crossing under more complex scenarios in which it may in fact be common (e.g. Trotter *et al*. 2014).

Valley crossing in asexual populations is increasingly well understood (Weissman *et al*. 2009). If the number of mutants produced every generation is high, as is the case in large populations with high mutation rates, a complex adaptation requiring multiple mutations that is destined to fix will eventually be generated de novo. This is sometimes called deterministic fixation of the multiple mutant (Weissman *et al*. 2009). In small populations, neutral or deleterious intermediate mutants generated by wild type individuals can drift to fixation. Further mutations necessary for the adaptation can then arise on intermediate mutant backgrounds and fix due to positive selection. This is the sequential fixation regime (Weissman *et al*. 2009). A third possibility occurs for intermediate population sizes where wild type individuals generate transient mutant subpopulations or “bubbles”. These bubbles are ordinarily doomed to extinction due to drift, but additional mutations in a lucky bubble may generate a complex adaptation that sweeps to fixation; this process has been referred to as “tunneling” (Iwasa *et al*. 2004; Weissman *et al*. 2009). On its own, valley crossing becomes more likely as the population size increases (Weissman *et al*. 2009). However, recent work suggests that valley crossing relative to the fixation of a simple beneficial mutation is least likely at intermediate population sizes where tunneling occurs Ochs and Desai (2015). This suggests that tunneling may be the most difficult mode of valley crossing. Recombination also affects the rate of valley crossing. Infrequent recombination decreases valley crossing times (Weissman *et al*. 2010). At high values of the recombination rate, however, individuals carrying the full complex adaptation outcross with the wild type and produce more deleterious intermediates that retard the growth of the multiple mutant. An additional effect that can heighten the rate at which valley crossing occurs is population subdivision (Bitbol and Schwab 2014): the population size within a subdivision is smaller, meaning that selection against potentially harmful intermediates is relaxed.

Previous studies of valley crossing focus primarily on populations where all individuals are equal in fitness except for the focal loci, i.e., the loci at which the individual mutations comprising the complex adaptation segregate. This is tantamount to assuming that, if there is variation in the background fitness of the population, it is negligible. In such populations, genetic drift governs the fate of neutral alleles, as well as the behavior of deleterious or beneficial alleles where they are rare (close to frequency zero) or common (close to frequency one). Alleles whose dynamics are primarily determined by genetic drift can be effectively modeled by a diffusion approximation (Wright 1945; Kimura 1955, 1957), and the ancestry of a population is described well by the classic Kingman coalescent (Kingman 1982) where pairs of branches in the genealogy merge back in time until the most recent common ancestor. When population sizes are very large or the fitness variation in the population is substantial, however, the behavior of genetic variation is governed more by genetic *draft* than by genetic drift (Gillespie 2000, 2001; Masel 2011; Neher and Shraiman 2011). Whereas drift is the effect of imperfect sampling from generation to generation, draft is the sum of hitchhiking effects due to selection on sites linked to the focal loci. Draft is fundamentally different from drift: for example, drafting populations experience large jumps in allele frequencies that cannot be encapsulated by a diffusion approximation (Neher and Shraiman 2011). They commonly have genealogies in which more than two branches merge at once and that are better described by alternative coalescent processes such as the Bolthausen-Sznitman coalescent (Neher and Hallatschek 2013; Brunet *et al*. 2007; Schweinsberg 2017). Additionally under genetic draft, the frequency spectrum of neutral alleles is non-monotonic, with a marked uptick near frequency one–or, equivalently, a depression in intermediate frequency alleles, which are quickly swept out of the population (Neher and Shraiman 2011; Kosheleva and Desai 2013; Neher and Hallatschek 2013). Only a handful of individuals near the “nose” of the fitness distribution are likely to persist, and they give rise to the bulk of the future population and carry linked alleles with them. The parameter that determines whether drift or draft is more important is the product of the population size *N* and the standard deviation in fitness *σ* (Neher and Hallatschek 2013).

It is increasingly realized that draft may play a critical role in shaping genetic diversity, especially in many microbial species, where population sizes can be large but “effective population sizes” are many orders of magnitude smaller (Masel 2011). The possibility that neutral variants are affected more strongly by selection at linked sites than by genetic drift, even in organisms like *Drosophila*, is one possible explanation for the “paradox of variation”, the fact that genetic diversity and population size often do not scale linearly (Gillespie 2000, 2001; Neher *et al.* 2013; Corbett-Detig *et al*. 2015). Draft likewise may have profound effects on the evolution of complex adaptations. Preliminary evidence for this comes from Neher and Shraiman (2011), who explored how draft affects valley crossing via stochastic tunneling by calculating the mutant bubble size distribution in a rapidly adapting population. They found that, compared to drift, draft generally causes the distribution of bubble sizes to drop off more quickly, meaning that small bubbles are more common but large bubbles are very rare. Thus, while draft may make valley crossing via tunneling more difficult, the total effect isn’t immediately obvious. Here, we extend these previous results by studying how genetic draft in asexual populations affects the rate of crossing fitness valleys across a range of population sizes.

The complicating factor is that, in an asexual population, the dynamics are governed almost entirely by what happens in the nose of the fitness distribution, and these dynamics can be highly stochastic. In previous approaches, these stochastic effects were smoothed due to the presence of recombination (Neher and Shraiman 2011, 2009). We therefore must focus primarily on simulation approaches, as analytical solutions for the behavior of a mutation in the nose of an asexual population are difficult to obtain. We show that fitness valley crossing occurs at an overall lower rate in rapidly adapting populations. However, this is consistent with the fact that *all* forms of adaptation are slowed down due to clonal interference and genetic background effects. In addition, we confirm that in rapidly adapting populations, adaptive fixations of alleles are more likely to be complex adaptations involving a fitness valley than they are in populations where genetic drift is the primary force shaping genetic variation at linked sites; essentially, genetic draft maintains the linkage that makes it possible to leap across fitness landscapes rather than adapt primarily by climbing to local peaks. These observations add to the growing intuition that complex adaptations that involve evolution across fitness valleys may not be a surprising result of the evolutionary process, but rather an expected one.

**Figure 1.**
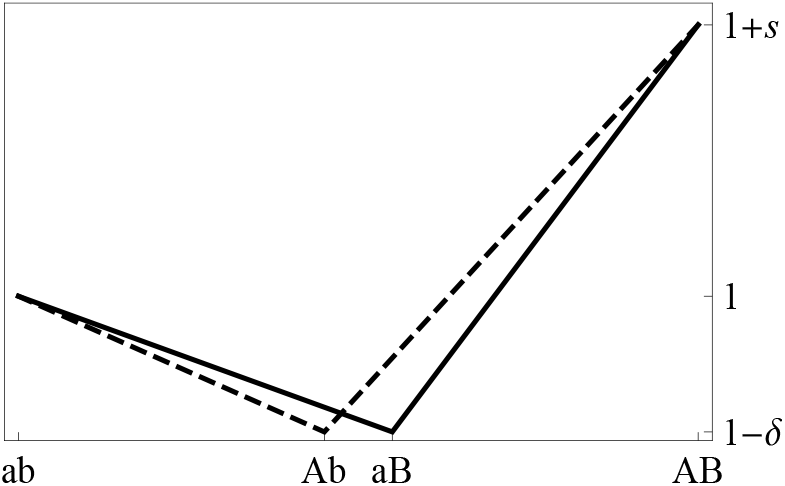
Fitness landscape

## Mathematical background

Our model features a population of *N* haploid individuals in which an effectively infinite number of “background” loci, with weak fitness effects, are currently segregating.

We focus on two loci, which are initially fixed for alleles *a* and *b*. These alleles mutate to *A* and *B* at rates *μ_A_* and *μ_B_*, respectively (we neglect back mutations). Genotypes *ab, Ab, aB*, and *AB* have fitnesses 1,1 — *δ_A_*, 1 — *δ_B_*, and 1 + *s* respectively, with *δ* ≥ 0 (intermediate genotypes are deleterious) and 1/*N* < *s* < 1 (double mutant *AB* is strongly beneficial); see 1. For the remainder of our analysis, we will assume that *μ_A_* = *μ_B_* = *μ* and *δ_A_* = *δ_B_* = *δ*. We further assume that the large number of loci of weak effect contribute to a constant background fitness variance *σ*^2^. By Fisher’s “fundamental theorem” of natural selection (Fisher 1930), the rate *υ* at which the mean fitness advances due to selection is set to *σ*^2^. If *x* is the background fitness of a lineage, then the distribution of background fitnesses *f(x)* is assumed to be roughly Gaus-sian: 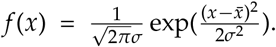. Given this background fitness variance, we are interested in the expected time 𝔼 [*T*] for the double mutant *AB* (our so-called “complex adaptation”) to arise and fix in the population. We emphasize that *σ*^2^ is only a *background* fitness variance and does not include the effects of the focal loci. For the most part, we assume that the focal loci are either at low enough frequency that they do not substantially affect the total fitness variance or that they are on the way to fixation and the population is destined to become roughly monomorphic (until new beneficial background mutations can be introduced).

In very large populations, enough double mutants are generated that the valley crossing time is dominated by the time required for a double mutant to sweep through the population. At the other extreme, in small populations, the crossing time is dominated by the wait for a deleterious single mutant to fix: only thereafter does a successful double mutant appear. We zero in on the intermediate case: stochastic tunneling, the most interesting and mathematically demanding mode of valley crossing. We will review existing theory for this case, discuss the difficulty in the mathematical analysis due to genetic draft, and then present results for valley crossing times using forward-time simulations across a range of population sizes that include the sequential fixation, stochastic tunneling, and semi-deterministic tunneling cases.

There are three components to the process of stochastic tunneling. First, a single mutant lineage must appear, which occurs with rate *N_μ_* If this lineage arises at time *t*_0_ and persists until time *T*, then it gives rise to 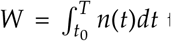 total individuals before going extinct, where *n*(*t*) is the number of mutant individuals extant at time *t*. As in the introduction, we refer to such a shortlived mutant lineage, and all the individuals generated within that lineage, as a mutant “bubble”: the time integrated number of individuals *W* is the “weight” of the bubble. During the lifetime of a bubble, the complex adaptation must appear and establish; that is, it must rise to a high enough frequency that fixation is almost guaranteed, which occurs at rate *μp*_fix_ where *μ*_fix_ is the probability that the double mutant reaches fixation from a single initial individual. If it establishes, the complex adaptation will quickly sweep to fixation. The first two steps can be folded in together and considered as one process, so that what is relevant is the expected time until a single mutant lineage arises that is destined to produce a successful double mutant. In this way,𝔼[*T*] =𝔼[*T*_0_ + *T*_1_], where *T*_0_ is the time to the first successful bubble-the first that produces a double mutant that is destined for fixation-and *T*_1_ is the time required for the double mutant to sweep. We assume that *T*_0_ ≫ *T*_1_, so that the wait time is dominated by the first term: the time required for *T*_1_ to establish will be of order 1/*s* (Desai and Fisher 2007), and the sweep time will be of order 1/*s* log(*Ns*), but *T*_0_ can be many orders of magnitude greater.

Most of the time, a bubble will simply arise and go extinct. If mutation and fixation are sufficiently rare events, then fixation of the double mutant can be modeled as a Poisson process: with probability 1 – *e*^−*μp*_*fix*_^*W*^^, the bubble with total size *W* will give rise to a successful double mutant. Computing the expected value of this quantity over all bubble sizes, 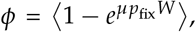, gives the probability that some bubble will lead to a successful valley crossing. One difficulty in computing the rate of tunneling in the high *N_σ_* case (where genetic draft is significant) is that the expected bubble size and fixation probability will both turn out to depend on the background fitness of an individual. That is to say, it is not possible to simply compute p_fix_ and the weight distribution by themselves, but rather one must convolute both of them over the background fitness. Individuals whose background fitness is near the nose of the fitness distribution will tend to give rise to longer-lived lineages, and if those individuals give rise to the complex adaptation, it will therefore likewise be more likely to fix. Thus, *W* will depend on the background fitness *x* and the valley depth *δ*, and *p*_fix_ will depend on *x* and the peak height s. These will have to be integrated over the fitness distribution *f*(*x*) in order to obtain the total crossing probability 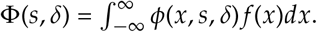. The expected valley crossing time then becomes 1/*N_μ_*Φ. An additional difficulty is that technically, *p*_fix_ will depend not on the fitness of the initial lineage in a bubble but on the fitness of the specific background on which the double mutant arose. These will not necessarily be the same; for example, individuals within the single mutant lineage may accumulate additional background mutations. Assuming that this does *not* happen, that is, assuming that *p*_fix_ depends on the fitness *x* of the founding single mutant lineage, is tantamount to assuming that bubbles are too short lived to accumulate background mutations that significantly affect their fitness.

We begin by considering the dynamics of *p*_fix_(*x, s*) before moving on to consider the distribution of bubble sizes. We focus on the case where background mutations are frequent but of weak effect-that is, the mean effect of background mutations e is small compared to *̄∊* but background mutations occur at a high rate *U_b_*. Hallatschek (2011) and Good and Desai (2014) showed that the fixation probability in this regime obeys the differential equation

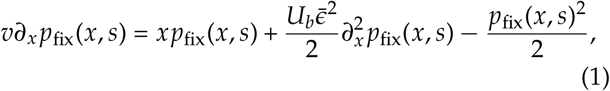

in which we have further assumed that *υ* ≈ *σ*^2^, which is tantamount to claiming that mutational effects are less important than selection in determining the advance of the mean fitness. For a neutral mutation (*s* = 0), the fixation probability is effectively zero at low *x* but increases exponentially up until *x_c_*, at which it becomes roughly linear in *x*. In this regime, 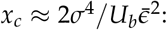 when *σ* is large, only mutations in the nose stand any chance of fixing unless they can be dragged to higher fitness by background mutations. The effect of introducing a nonzero *s* is essentially to shift the value of *x* and *xc*: for large negative *s*, *x_c_* may be far enough in the nose that it is unlikely any individuals will actually be present, and for large positive s, even unfit individuals can give rise to a lineage that is destined to fix.

In order to obtain the total valley crossing probability Φ, the only remaining element to compute is the distribution of mutant bubble sizes *W* as function of their background fitness *x*. Since the tunneling probability for a lineage with background fitness *x* is given by 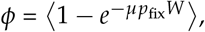, we do not need to compute the full distribution of *W* but rather only its Laplace transform: 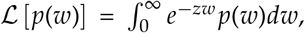 with *z* = *μp*_fix_. Neher and Shraiman (2011) showed that *ϕ* follows the equation

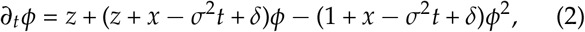

but their analysis also included a recombination term that made it possible to pull out the integrated Laplace transform 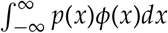 without the need to actually solve this equation. They also avoided the need to explicitly send time to infinity, which is necessary to incorporate all possible bubble sizes and lifetimes. Unfortunately this method is not available to us, and the nose dominance of the bubble size distribution, combined with the *x* dependence of *z* (via *z* = *μp*_fix_(*x*)), lead to difficulty in deriving a simple closed form expression for Φ. Therefore, we focus on simulation methods to study the dynamics of valley crossing.

## Simulation methods

To analyze our model, we perform forward simulations using a customized version of FFPopSim (Zanini and Neher 2012), a discrete time forward simulation package that implements a modified Wright-Fisher model. The simulation code is available via GitHub. In FFPopSim, individuals are organized by “clones” (sets of individuals with the same genome): each individual gives rise to a Poisson distributed number of offspring, with the mean dependent on the relative fitness. The mean offspring number is further adjusted to keep the population roughly at a pre-specified carrying capacity. We initialize a wild type population of *N* individuals consisting of haploid genomes of with *L* = 200 loci. Genetic loci are partitioned into two groups, the two “focal” loci where the epistatic alleles *A* and *B* will segregate and the remaining “background” loci that are responsible for the underlying fitness distribution. We set the focal loci to be at positions *L*/4 and 3*L*/4. For the background loci, we make use of a modified infinite sites model (Kimura 1971; Watterson 1975): any time a locus becomes monomorphic, a mutation at that locus is injected into a random individual in the population. In this way, the population experiences a constant influx of beneficial mutations at the background loci. These mutations’ fitness effects are drawn from an exponential distribution; the background fitness variance is set to a constant value *σ*^2^ by manually adjusting the selection coefficients every generation (they are multipled by the current variance and divided by *σ*^2^). In this way, the mutation rate *U_b_* and average fitness effect *̄∊* at the background loci are not parameters of our simulation model but must be directly measured from simulations; i.e., they are constrained by *L* and *σ*. We estimate them by manually counting the number of injected mutations every generation and averaging over the fitness effect at each locus.

Simulations are allowed to proceed for an equilibration time of *N*/10 generations to allow sufficient genetic diversity to be introduced. During this time, the mutation rate *μ* for the focal loci is set to zero. After equilibration, we set the per-site mutation rate *μ* for the focal loci (allowing forward and backward mutations) with single mutant fitness disadvantage *δ* and double mutant advantage s. When the double mutant reaches frequency 0.5, we consider it to have fixed. We then set the focal loci allele frequencies back to zero, randomize the selection coefficients for the genetic background by drawing anew from an exponential distribution, and allow the population to equilibrate for another *N*/10 generations. This ensures that independent valley crossing trials are independent despite occurring in the same simulation run.

We partitioned our simulations as follows. We first studied the effect of increasing *σ* in the sequential fixation, stochastic tunneling, semi-deterministic tunneling, and deterministic fixation regimes (Weissman *et al*. 2009). For small populations, sequential fixation applies when 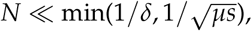, so that a successful mutant is unlikely to arise until the intermediate has fixed, but the intermediate’s fitness disadvantage is too weak for selection to inhibit it. Tunneling becomes relevant when *N* is too large for sequential fixation, i.e., the tunneling probability becomes more significant than the probability that the intermediate will fix: this mandates 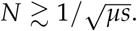 It also requires that *N* ≪ 1/*μ*, meaning that only one mutant lineage is likely to be extant at any given time and, if lineages do co-occur, they are unlikely to interfere with each other or substantially affect the mean fitness. By increasing *N* far above the 1/*μ* limit, multiple deleterious mutants are likely to occur every generation, reaching mutation-selection balance at low frequency and providing many opportunities for a lucky double mutant to appear; this is semi-deterministic tunneling. At still higher population sizes 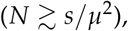 multiple double mutants are likely to appear every generation, and their dynamics can be modeled as deterministic.

Finally, we compared the dynamics of valley crossing versus sweeping mutations. We performed simulations with a sweeping focal beneficial mutation, with selection coefficient *s*_sweep_, at position *L*/2, with the background loci kept polymorphic as in the valley crossing simulations. We recorded the time for a beneficial mutation to arise and reach frequency 0.5, considering it to have fixed at that point. We then computed the ratio of the sweep time to the valley crossing time for a complex adaptation, with valley depth *δ* and larger fitness advantage *s*_valley_. To determine the effect of *σ* on this ratio, we performed simulations at both high and low *σ* and computed the ratio of these ratios: the result is a measure of the extent to which rapid adaptation increases the likelihood of valley crossing relative to the speed of simple sweeps.

### Data availability

Simulation and analysis scripts needed to replicate the results are available at https://github.com/tkessinger/draft_valley_crossing under the Creative Commons BY-NC-SA license (https://creativecommons.org/licenses/by-nc-sa/4.0/).

## Results

Here we present the key results of our simulation analysis. In general, we expect valley crossing to require more time in rapidly adapting populations (large *Nσ*) experiencing genetic draft; bubbles are doomed to extinction more quickly (lineages experience larger fluctuations in size and decline in frequency as the mean fitness advances) and beneficial lineages are less likely to fix, especially when valleys are shallow. Figures 2 and 3 confirm these expectations: the overall rate of valley crossing is generally slowed as *σ* increases. At higher *σ*, the effect of the genetic background in which an mutation finds it-self becomes more significant in determining the fate of that mutation than the fitness effect of the mutation itself. This genetic draft buffets the frequency of mutations so that these mutations behave as though they are more neutral than they really are: in effect, the fitness landscape is “flattened”. This depresses the fixation probability of the double mutant and other complex adaptations, which appears to account for the majority of the change in fixation time. In the case of stochastic tunneling, the overall size distribution of bubble sizes is reduced as well: the distribution *P*(*w*) scales not as *w*^−3/2^ but as *w*^−2^(Neher and Shraiman 2011), so there are fewer chances for the complex adaptation to arise on a mutant background and fix.

**Figure 2.**
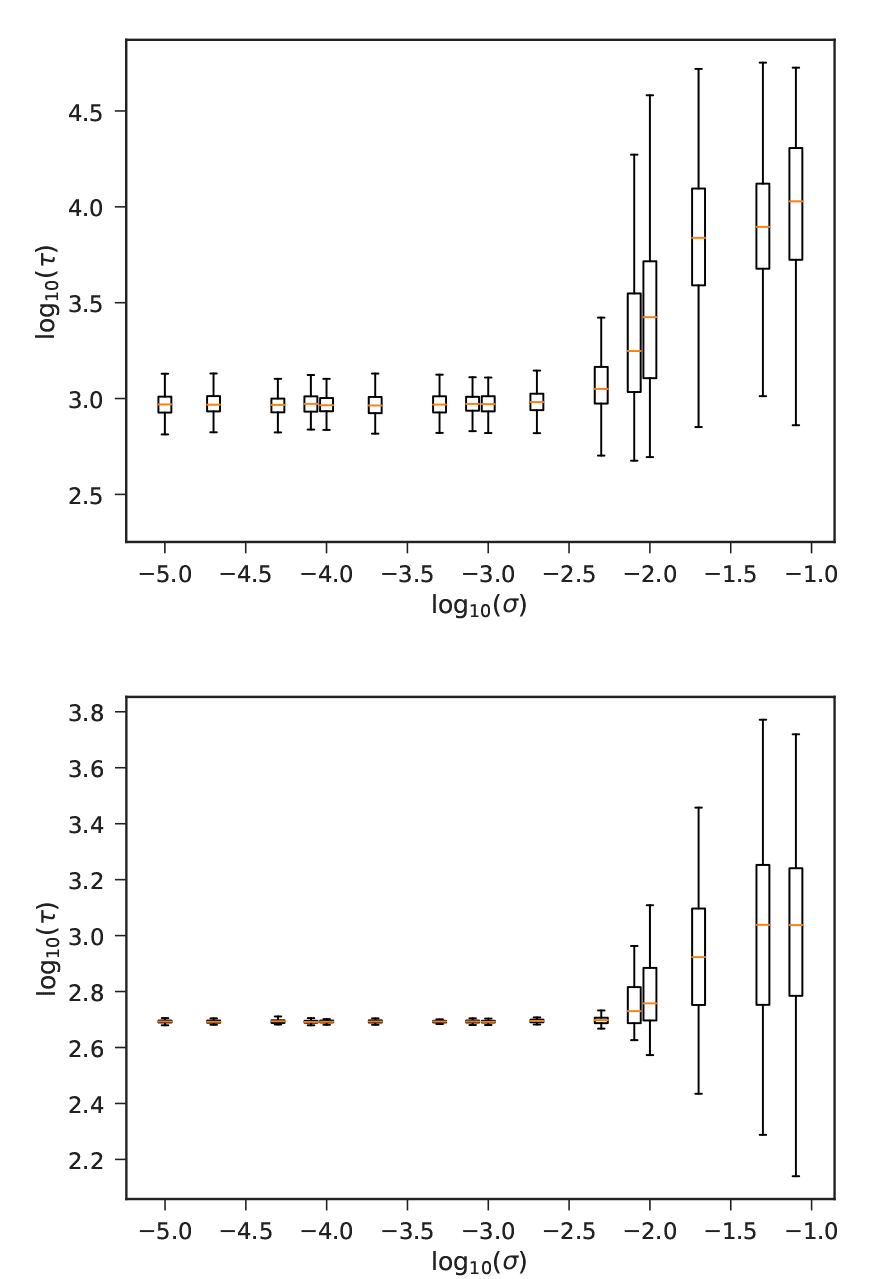
Valley crossing times in the semi-deterministic tunneling regime as a function of *σ*. In both figures, *s* = 10^−2^ and *δ* = 10^−3^: on the left, *N* = 10^5^ and *μ* = 10^−4^, and on the right, *N* = 10^6^ and *μ* = 10^−3^; the right figure has a high enough value of *Nμ*^2^ that fixation is almost deterministic. Orange lines are averages, boxes are interquartile ranges, and end caps are overall ranges. Each box represents about 100 simulation runs.

**Figure 3.**
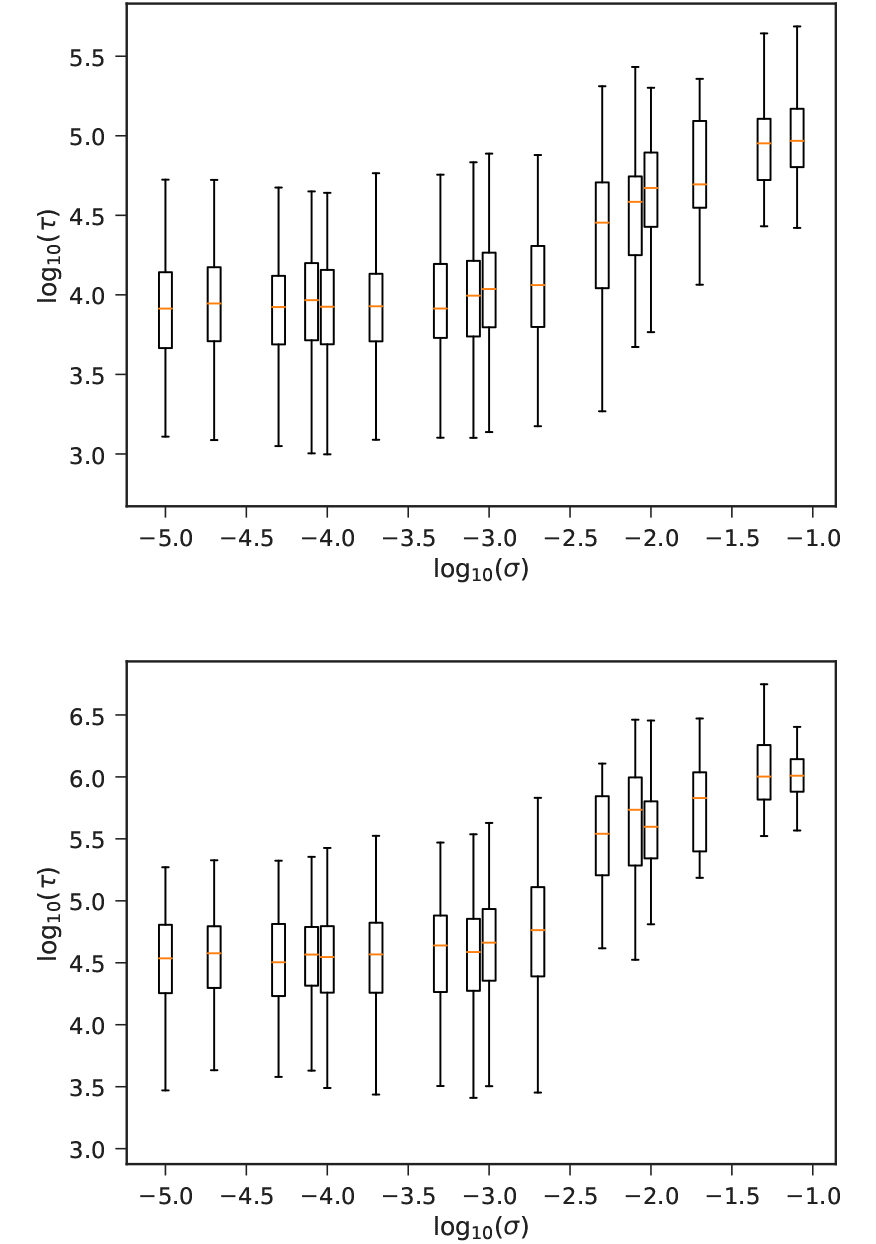
Valley crossing times in the sequential fixation (left) and tunneling (right) regimes as a function of *σ*. In both figures, *s* = 10^−2^ and *δ* = 10^−3^: on the left, *N* = 10^4^ and *μ* = 3 × 10^−5^, and on the right, *N* = 10^5^ and *μ* = 3 × 10^−6^. Labeling is as in figure 2.

On the other hand, the reduced effectiveness of selection should generally be helpful in crossing deeper valleys by mitigating the deleterious effect of single mutants. This can be seen in figure 4. At high values of *σ*, the fitness effect of the deleterious intermediate mutation has very little effect on the valley crossing time, which means that deep valleys can be crossed. This suggests that traversing rugged fitness landscapes by valley crossing may be enhanced in rapidly adapting populations as landscapes under genetic draft appear to be “flattened”.

So far, we can see that valley crossing is slower in rapidly adapting populations but the crossing time is less sensitive to the depth of the valley. How do these two effects combine to shape the likelihood of valley crossing in rapidly adapting populations experiencing genetic draft compared to populations only experiencing drift? To explore this, we first introduce the ratio *α* = *τ*_valley_/*τ*_sweep_, where *τ*_sweep_ is the time required for a single beneficial mutation to sweep to fixation and *τ*_valley_ is the time required for a valley crossing event to occur. By comparing how *α* changes as a function of the strength of selection on linked variation, *σ*, we can determine how *σ* affects the speed of valley crossing relative to simple beneficial sweeps. Higher (or lower) values of *α* mean that valley crossing times increase (or decrease) relative to the time required for a simple beneficial sweep. Though the precise value of this ratio depends on *δ* and on the selection coefficients *s*_valley_ and *s*_sweep_ of the complex adaptation and the single beneficial mutation, respectively, a general trend nonetheless emerges. In Figure 5, we set *s*_sweep_ = 0.01, *s*_sweep_ = 0.1, and *μ* = 10^−5^. Figure 5b shows the expected pattern that deeper valleys in smaller populations take longer to cross. At a higher value of *σ* valley depth has a weaker effect on the valley crossing time in both an absolute sense (Figure 5a) and relative to the time for a single beneficial sweep (Figure 5b). Moreover, values of *α* are generally lower for the population with higher *σ* experiencing genetic draft than for the population with lower *σ* experiencing only drift. This implies that draft tends reduce the valley crossing time relative to the time for a single beneficial mutation to sweep.

**Figure 4.**
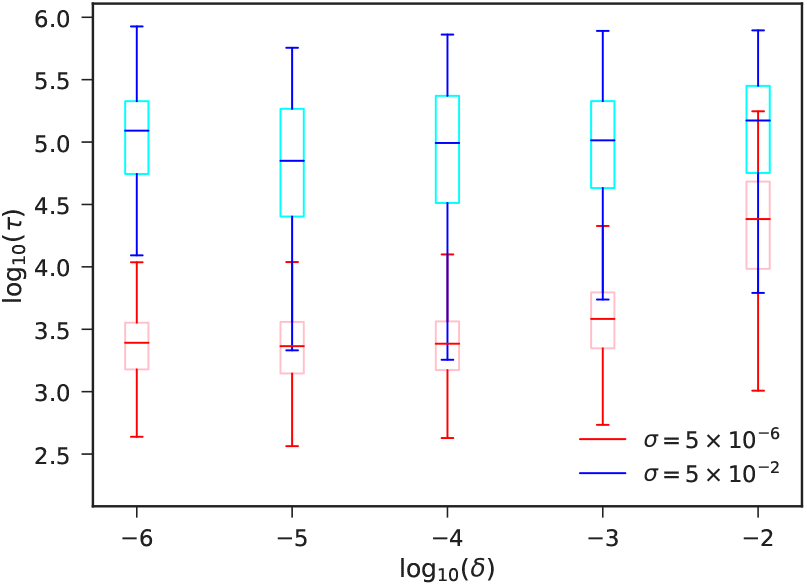
Valley crossing times for varied *σ* and *δ*. In these simulations, *N* = 10^5^, *μ* = 3 × 10^−6^, and *s* = 10^−2^. Note the lack of dependence on *δ* at high *σ*.

We can get a clearer picture of how draft affects valley crossing times relative to single beneficial sweeps by considering the ratio *α_σ_2__*/*α_σ_1__* where *σ*_1_ = 10^−6^ and *σ*_2_ = 10^−2^, which is a measure of the extent to which rapid adaptation speeds up valley crossing relative to sweeping (lower values indicate that rapid adaptation favors valley crossing). The ratio can be seen in Figure 6. Overall, we find that raising *σ* increases the extent to which valley crossing is favored for lower population sizes: for this combination of parameters, the condition is *N* < 10^5^. In this regime, raising *δ* also increases the degree to which valley crossing is favored under draft, which means that the deeper the valley, the more that draft increases the likelihood of valley crossing relative to a single beneficial mutation. The exception is at very high values of *N* and *δ* and low values of μ, where draft turns out again not to be helpful; see the lower right part of the leftmost plot. In this regime, *δ* is on the order of *σ* in our draft simulations, meaning that single mutant lineages are rapidly cut off by the advancing mean fitness and are likely almost never present in the nose of the fitness distribution. Essentially, the population size is large enough that intermediate mutants can be “seen” by selection, but the fitness detriment is severe enough compared to *σ* that draft is no longer helpful in allowing mutant lineages to reach high frequency. This problem disappears as *μ* increases (center and right plots) and the population is beyond the deleterious tunneling barrier where the valley crossing time depends on the depth of the valley δ.

**Figure 5.**
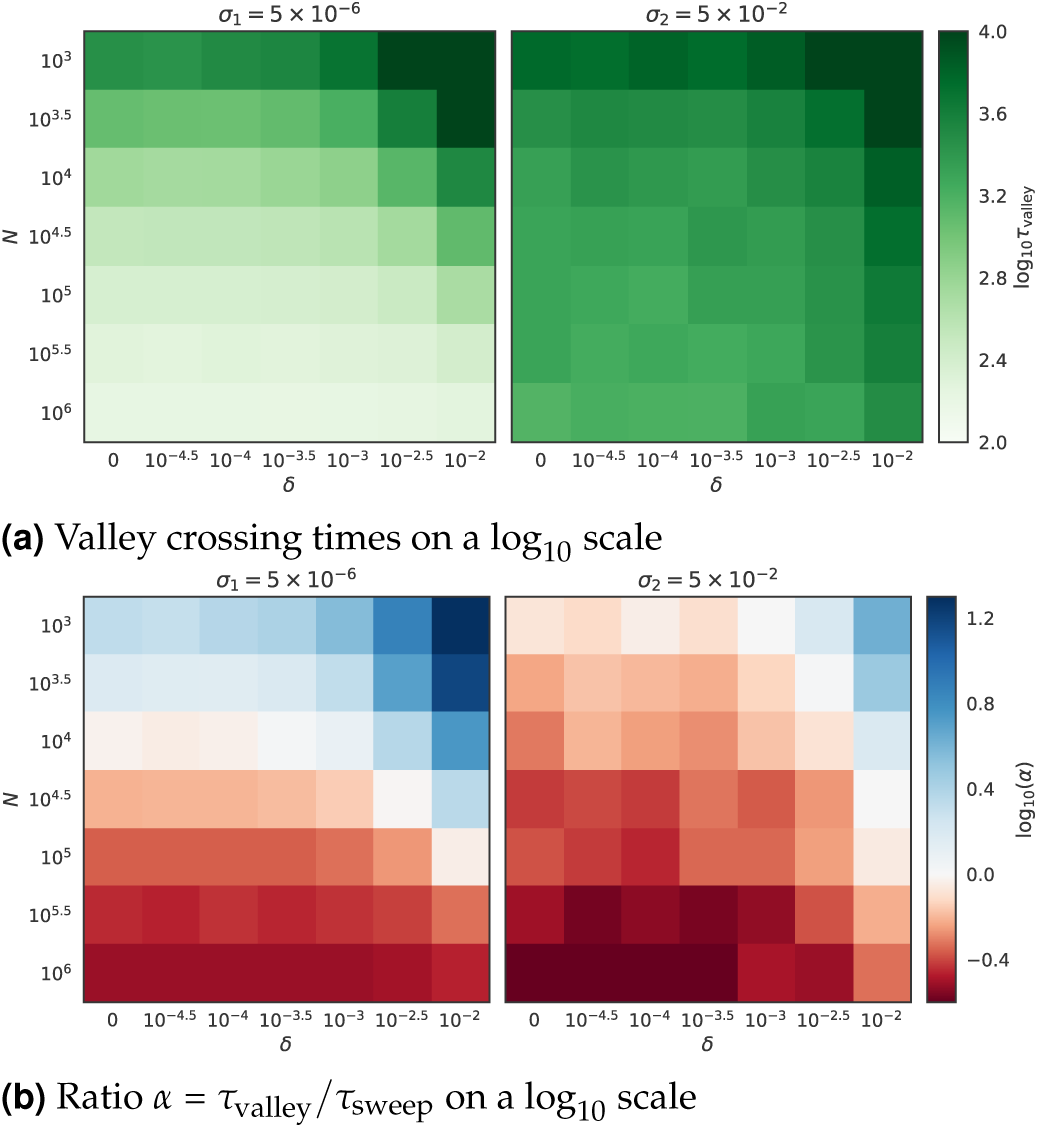
Behavior of valley crossing times for varied *δ*, N, and *σ*. In all cases, *s*_sweep_ = 0.01, *s*_valley_ =0.1, and *μ* = 10^−5^. Panel (a) shows that valley crossing times *τ* increase with *δ* and *σ* but decrease with *N*, as predicted. Panel (b) shows the ratio *α* = *τ*_valley_ /*τ*_sweep_ as a function of *δ*, *N*, and *σ*.

## Discussion

We have used simulation methods to examine the rate of crossing fitness valleys in rapidly adapting populations that experience genetic draft. We find that although the overall valley crossing time is longer in rapidly adapting than in slowly adapting (drifting) populations, a higher fraction of adaptive events in rapidly adapting populations will occur through valley crossing than in slowly adapting populations. This mean that the genetic draft that results from rapid adaptation accelerates the rate at which populations escape from local peaks in a fitness landscape and move towards global peaks. Thus, adaptation at loci unhindered by genetic draft occurs faster but is more likely to get stuck on a local peak in a rugged fitness landscape.

The source of this difference in behavior between populations experiencing genetic draft and those experiencing drift comes from two effects. First, the effect of the deleterious intermediate in the rapidly adapting population is much smaller: for 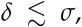 the precise value of *δ* almost does not affect the crossing rate. Second, the time to the evolution of even *simple* adaptations (i.e., ones requring substitution only at a single locus) is much longer in rapidly adapting populations. This means that the presence of a fitness valley is not as steep of an impediment to the evolution of a complex adaptation for a drafting population as it is for a drifting one.

**Figure 6.**
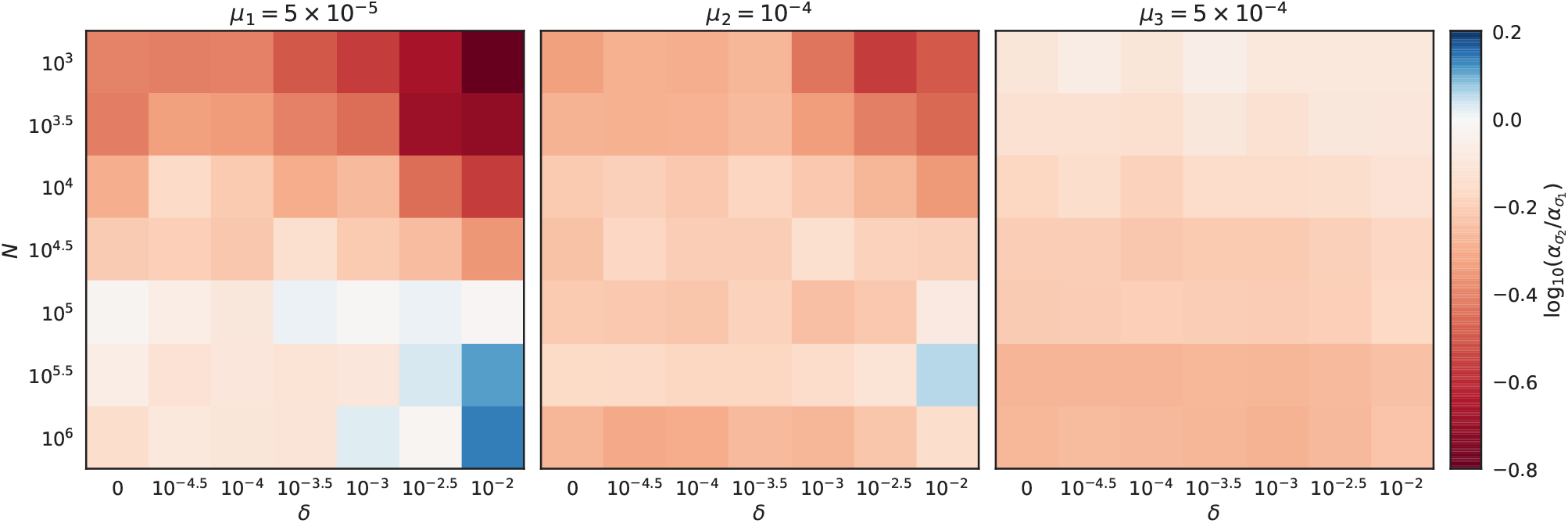
Ratio of *α* values at high and low *σ*(log(*α*_*σ*_2__/*α*_*σ*_1__ where *σ*_1_ = 10^−6^ and *σ*_2_ = 10^−2^), for varied values of *μ*. Genetic draft is helpful for crossing deep valleys in the sequential fixation and tunneling regimes, though at large population sizes it can slow down the crossing rate for deep valleys; at higher values of *μ* this limitation disappears.

Our results complement and build on those of Ochs and Desai (2015), who compared valley crossing and a sweeping beneficial mutation in a slowly evolving population. They considered the scenario where the sweeping beneficial mutation competes with a particular complex adaptation, whereas we treat these two scenarios separately. They find that large populations are more likely to cross valleys than fix a simple beneficial mutation than small populations (for comparison, see the *σ* = 5 × 10^−6^ panel in our Figure 5b). However, they also find that intermediate-sized populations in which stochastic tunneling occurs are the least likely to cross a fitness valley instead of fixing a beneficial mutation. In contrast, our results suggest that the stochastic tunneling regime is where rapid adaptation and genetic draft increase the likelihood of valley crossing relative to a simple beneficial mutation. Thus, it is possible that genetic draft improves the likelihood of valley crossing precisely where it is least likely under genetic drift.

We have focused exclusively on asexual populations, but extensions to sexual populations are possible. The relevant parameter is likely to be the product of *N* and the proportion of the fitness standard deviation segregating in a small, effectively asexual block, *σ_b_* (Neher *et al*. 2013). If both *Nσ* and *Nσ_b_* are large, and the focal loci segregate in the same block, then the analysis presented here should determine the crossing time. If, on the other hand, *Nσ* is large but *Nσ_b_* is small, or the map distance between the focal loci is much larger than the block length, then recombination between the focal loci can be modeled as “free”, with individuals effectively shuffling their entire genomes every generation. In that case, the communal recombination analysis of Neher and Shraiman (2011) and Neher *et al*. (2010) is likely to be important: the relationship between *σ* and the recombination rate *ρ* determines the crossing rate. If *σ* is large, then draft dominates the crossing rate: if *ρ* is large, then drift dominates it.

One common feature of the effect of genetic draft and population subdivision on valley crossing is that they both increase the appearance of complex adaptations by reducing selection against deleterious intermediates (i.e., the valley genotypes). In contrast, high migration rates and high recombination rates (Neher and Shraiman 2009) allow selection to more easily remove deleterious intermediates. This commonality has implications for Wright’s “shifting balance theory” (Wright 1932), which posits that population subdivision can enhance evolution on rugged fitness landscapes: deleterious intermediates accumulate due to genetic drift induced by subdivision, and subsequent beneficial mutations fix both in locally and are spread globally due to migration. However, it has long been known that the shifting balance theory has important limitations due to the fact that too much migration prevents the accumulation of deleterious intermediates and too little reduces the success of genotypes from the fitter peaks and prevents them from spreading throughout the metapopulation (Coyne *et al*. 2000; Van Cleve and Weissman 2015). Recent work by Bitbol and Schwab (2014) confirms the fact that intermediate migration rates yield the fastest valley crossing times; however, it also finds that valley crossing is faster in subdivided populations with these intermediate migration rates than in panmictic populations.

In light of our results that show that genetic draft can improve the likelihood of valley crossing, populations which experience both subdivision *and* rapid adaptation are ideal candidates in which the shifting balance theory may apply. Such populations include HIV, in which host-specific adaptations evolve quickly (Zhang *et al*. 1997; Wain *et al*. 2007; Dapp *et al*. 2017; Theys *et al*. 2018) and in which deleterious mutations are known to hitchhike to high frequency (Zanini and Neher 2013; *Zanini et al*. 2015). Our work also shines light on the likely pathway through which multi-drug resistance evolves in HIV. In recent years of the HIV pandemic, resistance has generally been slower to evolve (Feder *et al*. 2015) because more and stronger drugs are used. There are two possible explanations for this. One is that the use of anti-HIV drugs decreases the number of virions segregating within an individual, thus lowering the number of possible chances for a resistant phenotype to appear. The other is that such drugs, especially when used in concert, contort the fitness landscape so that evolution of a phenotype that is resistant to a drug cocktail is more difficult. Both factors are likely to play a role. However, since HIV is a population in which draft, not drift, is the dominant stochastic force and hence population size is less important in determining the evolutionary dynamics, our work suggests that the second factor is likely to be more significant in HIV compared to organisms in which draft is less important.

Our results for the dynamics of valley crossing under genetic draft are likely to be important for other evolutionary problems in which stochastic forces are known to be important. For example, there is a sort of duality between the evolution of complex adaptations and the evolution of cooperation, which can be seen as a complex behavioral adaptation where the highest per capita fitness at equilibrium requires the combination of multiple cooperative individuals (i.e., the combination of genes among different individuals instead among different loci). In social evolution theory, high migration rates between demes, which cause organisms to be less likely to interact with close kin, disfavor the evolution of cooperation: likewise, when migration rates are low, cooperation can be favored (Hamilton 1970; Rousset 2004; Van Cleve 2015). Evolutionary forces which strengthen existing associations between loci are more likely to lead to such complex phenotypes: evolutionary forces which weaken these associations hinder their evolution. This reasoning applies whether the loci in question appear in the same individual (as in the case of sign epistasis) or in different individuals (as in the case of social evolution). These effects can even interact: cooperation can function as an additional evolutionary force that favors valley crossing (Obolski *et al*. 2017) and can in some cases be a valley crossing adaptation in itself (Van Cleve and Lehmann 2013). The interplay between cooperation and valley crossing is an area that needs further study, as it may shed light on the evolution of cooperation within large microbial populations such as yeast (Gore *et al*. 2009; Sanchez and Gore 2013) and bacterial biofilms (Rainey and Rainey 2003; van Gestel *et al*. 2014).

## Acknowledgments

This work is supported by the National Science Foundation under Cooperative Agreement No. 1355438.

## Literature Cited

Aguilar-Rodríguez, J., J. L. Payne, and A. Wagner, 2017 A thousand empirical adaptive landscapes and their navigability. Nat. Ecol. Evol. 1: 0045.

Bitbol, A.-F. and D. J. Schwab, 2014 Quantifying the role of population subdivision in evolution on rugged fitness landscapes. PLoS Comput. Biol. 10: e1003778.

Brunet, E., B. Derrida, A. H. Mueller, and S. Munier, 2007 Effect of selection on ancestry: An exactly soluble case and its phenomenological generalization. Phys. Rev. E 76: 041104.

Corbett-Detig, R. B., D. L. Hartl, and T. B. Sackton, 2015 Natural selection constrains neutral diversity across a wide range of species. PLoS Biol. 13: 1–25.

Coyne, J. A., N. H. Barton, and M. Turelli, 2000 Is wright’s shifting balance process important in evolution? Evolution 54: 306–317.

Dapp, M. J., K. M. Kober, L. Chen, D. H. Westfall, K. Wong, et al., 2017 Patterns and rates of viral evolution in hiv-1 subtype b infected females and males. PLOS ONE 12: 1–30.

Darwin, C., 1859 On the Origin of Species by Means of Natural Selection. John Murray, London.

de Visser, J. A. G. M. and J. Krug, 2014 Empirical fitness landscapes and the predictability of evolution. Nat. Rev. Genet. 15: 480–490.

Dercole, F. and S. Rinaldi, 2008 Analysis of Evolutionary Processes: The Adaptive Dynamics Approach and Its Applications. Princeton University Press, Princeton, N.J.

Desai, M. M. and D. S. Fisher, 2007 Beneficial mutation–selection balance and the effect of linkage on positive selection. Genetics 176: 1759–1798.

Feder, A. F., S.-Y. Rhee, R. W. Shafer, D. A. Petrov, and P. S. Pennings, 2015 More efficacious drugs lead to harder selective sweeps in the evolution of drug resistance in hiv-1. bioRxiv.

Fisher, R. A., 1930 The Genetical Theory of Natural Selection. The Clarendon Press, Oxford.

Geritz, S. A. H., E. Kisdi, G. Meszena, and J. A. J. Metz, 1998 Evolutionarily singular strategies and the adaptive growth and branching of the evolutionary tree. Evol. Ecol. 12: 35–57.

Gillespie, J. H., 1983 A simple stochastic gene substitution model. Theor. Popul. Biol. 23: 202–215.

Gillespie, J. H., 1991 The Causes of Molecular Evolution. Oxford University Press, New York.

Gillespie, J. H., 2000 Genetic drift in an infinite population: The pseudohitchhiking model. Genetics 155: 909–919.

Gillespie, J. H., 2001 Is the population size of a species relevant to its evolution? Evolution 55: 2161–2169.

Good, B. H. and M. M. Desai, 2014 Deleterious passengers in adapting populations. Genetics 198: 1183–1208.

Gore, J., H. Youk, and A. van Oudenaarden, 2009 Snowdrift game dynamics and facultative cheating in yeast. Nature 459: 253 EP –.

Hallatschek, O., 2011 The noisy edge of traveling waves. Proc. Natl. Acad. Sci. U. S. A. 108: 1783–7.

Hamilton, W. D., 1970 Selfish and spiteful behaviour in an evolutionary model. Nature 228: 1218–1220.

Iwasa, Y., F. Michor, and M. A. Nowak, 2004 Stochastic tunnels in evolutionary dynamics. Genetics 166: 1571–1579.

Kimura, M., 1955 Solution of a process of random genetic drift with a continuous model. Proc. Natl. Acad. Sci. U. S. A. 41: 144–150.

Kimura, M., 1957 Some problems of stochastic processes in genetics. Ann. Math. Stat. 28: 882–901.

Kimura, M., 1971 Theoretical foundation of population genetics at the molecular level. Theor. Popul. Biol. 2: 174–208.

Kingman, J. F. C., 1982 The coalescent. Stochastic Process. Appl. 13: 235–248.

Kosheleva, K. and M. M. Desai, 2013 The dynamics of genetic draft in rapidly adapting populations. Genetics 195: 1007–1025.

Masel, J., 2011 Genetic drift. Curr. Biol. 21: R837–R838.

Neher, R. A. and O. Hallatschek, 2013 Genealogies of rapidly adapting populations. Proc. Natl. Acad. Sci. U. S. A. 110: 437–442.

Neher, R. A., T. A. Kessinger, and B. I. Shraiman, 2013 Coalescence and genetic diversity in sexual populations under selection. Proc. Natl. Acad. Sci. U. S. A. 110: 15836–15841.

Neher, R. A. and B. I. Shraiman, 2009 Competition between recombination and epistasis can cause a transition from allele to genotype selection. Proc. Natl. Acad. Sci. U. S. A. 106: 6866–6871.

Neher, R. A. and B. I. Shraiman, 2011 Genetic draft and quasi-neutrality in large facultatively sexual populations. Genetics 188: 975–996.

Neher, R. A., B. I. Shraiman, and D. S. Fisher, 2010 Rate of adaptation in large sexual populations. Genetics 184: 467–481.

Obolski, U., O. Lewin-Epstein, E. Even-Tov, Y. Ram, and L. Hadany, 2017 With a little help from my friends: cooperation can accelerate the rate of adaptive valley crossing. BMC Evol. Biol. 17: 143.

Obolski, U., Y. Ram, and L. Hadany, 2018 Key issues review: evolution on rugged adaptive landscapes. Rep. Progr. Phys. 81: 012602.

Ochs, I. E. and M. M. Desai, 2015 The competition between simple and complex evolutionary trajectories in asexual populations. BMC Evol. Biol. 15: 55.

Orr, H. A., 1998 The population genetics of adaptation: The distribution of factors fixed during adaptive evolution. Evolution 52: 935–949.

Rainey, P. B. and K. Rainey, 2003 Evolution of cooperation and conflict in experimental bacterial populations. Nature 425: 72 EP –.

Rousset, F., 2004 Genetic Structure and Selection in Subdivided Populations, volume 40. Princeton University Press, Princeton, N.J.

Sanchez, A. and J. Gore, 2013 Feedback between population and evolutionary dynamics determines the fate of social microbial populations. PLoS Biol. 11: 1–9.

Schweinsberg, J., 2017 Rigorous results for a population model with selection ii: genealogy of the population. Electron. J. Probab. 22: 54 pp.

Szendro, I. G., M. F. Schenk, J. Franke, J. Krug, and J. A. G. M. d. Visser, 2013 Quantitative analyses of empirical fitness landscapes. J. Stat. Mech. Theory Exp. 2013: P01005.

Theys, K., P. Libin, A.-C. Pineda-Peña, A. Nowé, A.-M. Vandamme, et al., 2018 The impact of HIV-1 within-host evolution on transmission dynamics. Current Opinion in Virology 28: 92 – 101, Emerging viruses: intraspecies transmission Viral Immunology.

Trotter, M. V., D. B. Weissman, G. I. Peterson, K. M. Peck, and J. Masel, 2014 Cryptic genetic variation can make irreducible complexity a common mode of adaptation in sexual populations. Evolution 68: 3357–3367.

Van Cleve, J., 2015 Social evolution and genetic interactions in the short and long term. Theor. Popul. Biol. 103: 2–26.

Van Cleve, J. and L. Lehmann, 2013 Stochastic stability and the evolution of coordination in spatially structured populations. Theor. Popul. Biol. 89: 75 – 87.

Van Cleve, J. and D. B. Weissman, 2015 Measuring ruggedness in fitness landscapes. Proc. Natl. Acad. Sci. U. S. A. 112: 7345–7346.

van Gestel, J., F. J. Weissing, O. P. Kuipers, and A. T. Kovács, 2014 Density of founder cells affects spatial pattern formation and cooperation in bacillus subtilis biofilms. ISME J. 8: 2069–2079.

Wain, L. V., E. Bailes, F. Bibollet-Ruche, J. M. Decker, B. F. Keele, et al., 2007 Adaptation of hiv-1 to its human host. Mol. Biol. Evol. 24: 1853–1860.

Watterson, G. A., 1975 On the number of segregating sites in genetical models without recombination. Theor. Popul. Biol. 7: 256–76.

Weissman, D. B., M. M. Desai, D. S. Fisher, and M. W. Feldman, 2009 The rate at which asexual populations cross fitness valleys. Theor. Popul. Biol. 75: 286 – 300, Sam Karlin: Special Issue.

Weissman, D. B., M. W. Feldman, and D. S. Fisher, 2010 The rate of fitness-valley crossing in sexual populations. Genetics 186: 1389–1410.

Wright, S., 1932 The roles of mutation, inbreeding, cross-breeding and selection in evolution. In Proceedings ofthe Sixth International Congress on Genetics, volume 1, pp. 356–366.

Wright, S., 1945 The differential equation of the distribution of gene frequencies. Proc. Natl. Acad. Sci. U. S. A. 31: 382–389.

Zanini, F., J. Brodin, L. Thebo, C. Lanz, G. Bratt, et al., 2015 Population genomics of intrapatient hiv-1 evolution. eLife 4: e11282.

Zanini, F. and R. A. Neher, 2012 Ffpopsim: an efficient forward simulation package for the evolution of large populations. Bioinformatics 28: 3332–3333.

Zanini, F. and R. A. Neher, 2013 Quantifying selection against synonymous mutations in hiv-1 env evolution. J. Virol. 87: 11843–11850.

Zhang, L., R. S. Diaz, D. D. Ho, J. W. Mosley, M. P. Busch, et al., 1997 Host-specific driving force in human immunodeficiency virus type 1 evolution in vivo. J. Virol. 71: 2555–61.

